# Impact of stimulus size and orientation on individual face decoding in monkey face-selective cortex

**DOI:** 10.1101/243154

**Authors:** Jessica Taubert, Goedele Van Belle, Rufin Vogels, Bruno Rossion

## Abstract

Face-selective neurons in the monkey temporal cortex discharge at different rates in response to pictures of different faces. Here we tested whether the population response of neurons in the face-selective area ML (located in the middle Superior Temporal Sulcus) tolerates two affine transformations; one, picture-plane inversion, known to have a deleterious impact on the average response of face-selective neurons and the other, stimulus size, thought to have little or no impact on face-selective neurons. We recorded the response of 57 ML neurons in two monkeys. Face stimuli were presented at two sizes (10 and 5 degrees of visual angle) and two orientations (upright and inverted). The results indicate that different faces elicited distinct patterns of activity across ML neurons that were tolerant of changes in size. However, the results of the orientation manipulation were mixed; despite observing a reduced response to inverted faces, classifier performance was above chance for both upright and inverted faces and the classification score did not differ significantly for inverted and upright faces. We conclude that population responses in area ML to different faces are dependent on stimulus orientation but are more tolerant to changes in stimulus size.

## Introduction

Single neurons that respond selectively to face compared to non-face visual stimuli were identified in the inferior temporal (IT) cortex of non-human primates over forty years ago^1^. Face-selective neurons in the IT cortex of macaque monkeys are characterized by their high category-selectivity (i.e., responding at least twice as much to faces than other similar shapes and objects^2–4^) across scale and position changes of the retinal image^5–7^. Most face-selective neurons respond with different orders of magnitude, i.e. firing rate, to pictures of different individual faces ^2,3,8–10^ offering a potential mechanism for achieving individual face discrimination ^11–13^

fMRI studies have defined a cortical face processing system in the monkey brain comprised of multiple interconnected, functionally-defined regions or ‘patches’ ^3,5,9,14–20^ in the last 10 years researchers have been able to use these fMRI maps to guide single cell recordings in monkeys, in order to understand the role of each functionally-defined patch, and to shed light on how face representations are successively transformed along the ventral visual pathway ^3,8,9,13^.

Among the face-selective clusters identified in the monkey brain, area ML, in the middle lateral section of the Superior Temporal Sulcus (STS) is the most consistently observed and investigated^3,18^. Using population decoding, studies have shown that activity in area ML, combined with MF (a face-selective patch in the fundus of the middle STS region), is identity-selective, although this decoding performance is thought to be limited by head orientation^3,8,9,13,21,22^.

Based on these observations, the aim of the present study is to test whether the identity of 12 randomly selected faces can be decoded from single neuron output in area ML, and, most importantly, the extent to which this performance depends on stimulus size and orientation, i.e., picture-plane inversion. A number of studies combining functional imaging and single cell recordings have reported that the average firing rate of ML neurons is systematically lower when face stimuli are turned upside down^3,5,21^. However, none of these studies have tested whether the unique patterns of responses across ML neurons elicited by different face identities are affected by such changes in orientation.

## Results

### Analysis of average activity across neurons

Fifty-seven face-selective neurons were recorded in area ML across two monkeys (average Face Selectivity Index (*FSI;* see Methods) = 0.67, sd = 0.25). Average normalized firing rate was analyzed using a 2 (*Size*) x 2 (*Orientation*) x 12 (*Face identity*) repeated measures ANOVA corrected for violations of sphericity using the Greenhouse-Geisser method. Neurons in ML responded stronger to upright than inverted faces (main effect of *Orientation*, F(1,56) = 13.19, *p* < 0.001), which extends previous findings in this region ^3,5^ and shows that the preference for upright faces remains even when averaging across multiple identities and stimulus sizes (Figure 2A). There was also a main effect of *Face Identity*, indicating that mean response strength was greater for some identities than others (F(8.26,462.49) = 2.67, *p* = 0.006). In contrast, the *Size* of the face stimulus did not change the response strength of ML neurons significantly (F(1,56) = 3.33, *p* = 0.08; see Figure 2B), and there was no significant interaction between *Size* and *Face identity* (F(8.43,472.39) = 1.43, *p* = 0.17). When averaging across face identities, there was no interaction either between *Orientation* and *Size* (F(1,56) = 0.24, *p* = 0.62) and the three way interaction also failed to reach significance (F(8.43,471.90) = 1.86, *p* = 0.06). For individual monkey data, please see Figure 2C.

**Figure 1:**
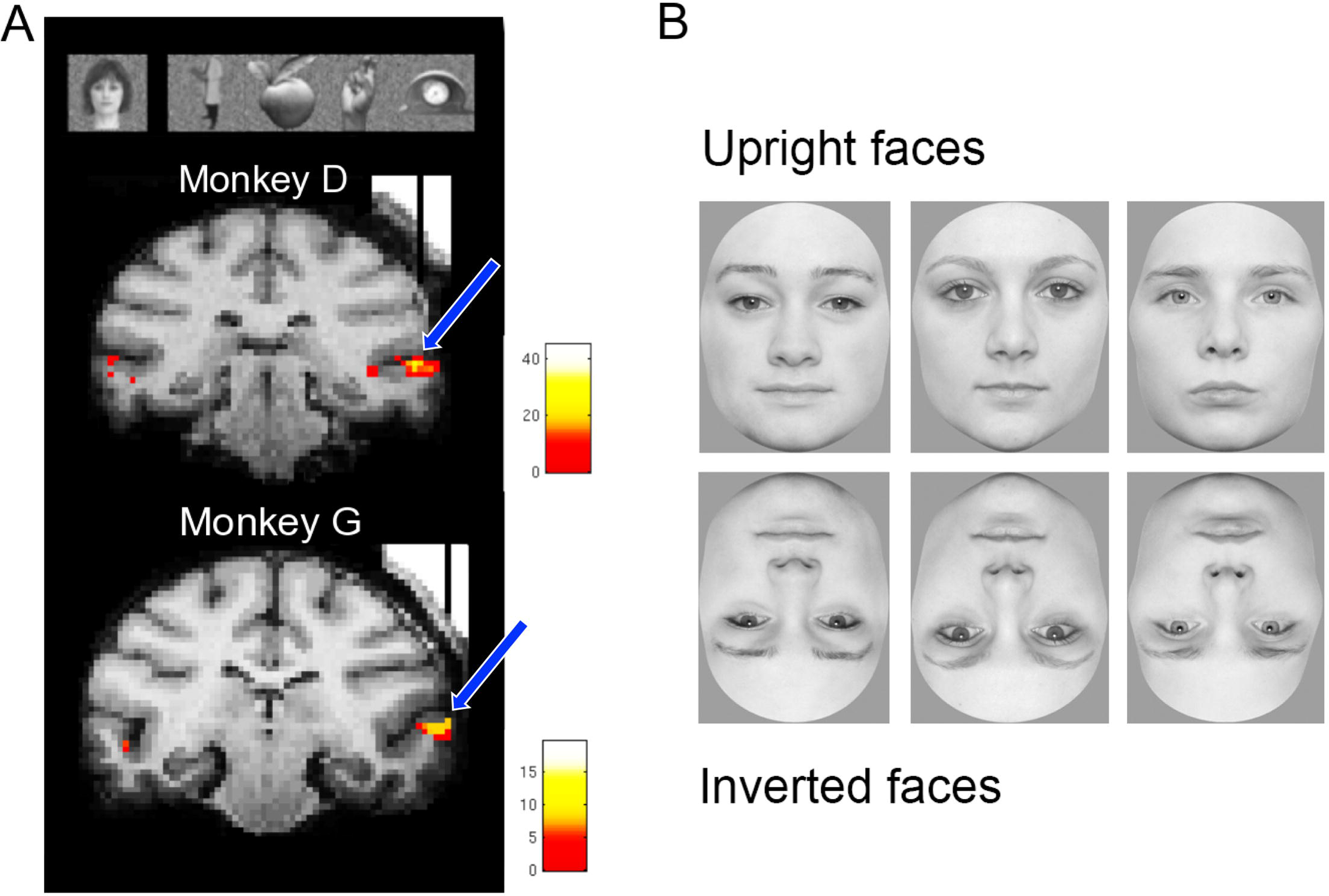
FUNCTIONAL MAPS AND EXPERIMENTAL STIMULI (A) The Area ML was defined in both monkeys using an fMRI block-design localizer with 5 categories of objects (the contrast was defined as [faces] – [bodies, fruit, hands, and gadgets]). In both cases, MION activation is superimposed on a high resolution anatomical scan obtained with tungsten markers positioned in the recording chamber grid to indicate the recording position. The t-maps are thresholded at p<0.05 (Family-Wise Error), corresponding to a t >4.9. The recording location in both monkeys is indicated by both the vertical position of the tungsten markers and blue arrows that have been superimposed on the scans. (B) Illustrative examples of the individual face pictures used (NB here we show examples of the orientation manipulation and not the size manipulation).

**Figure 2:**
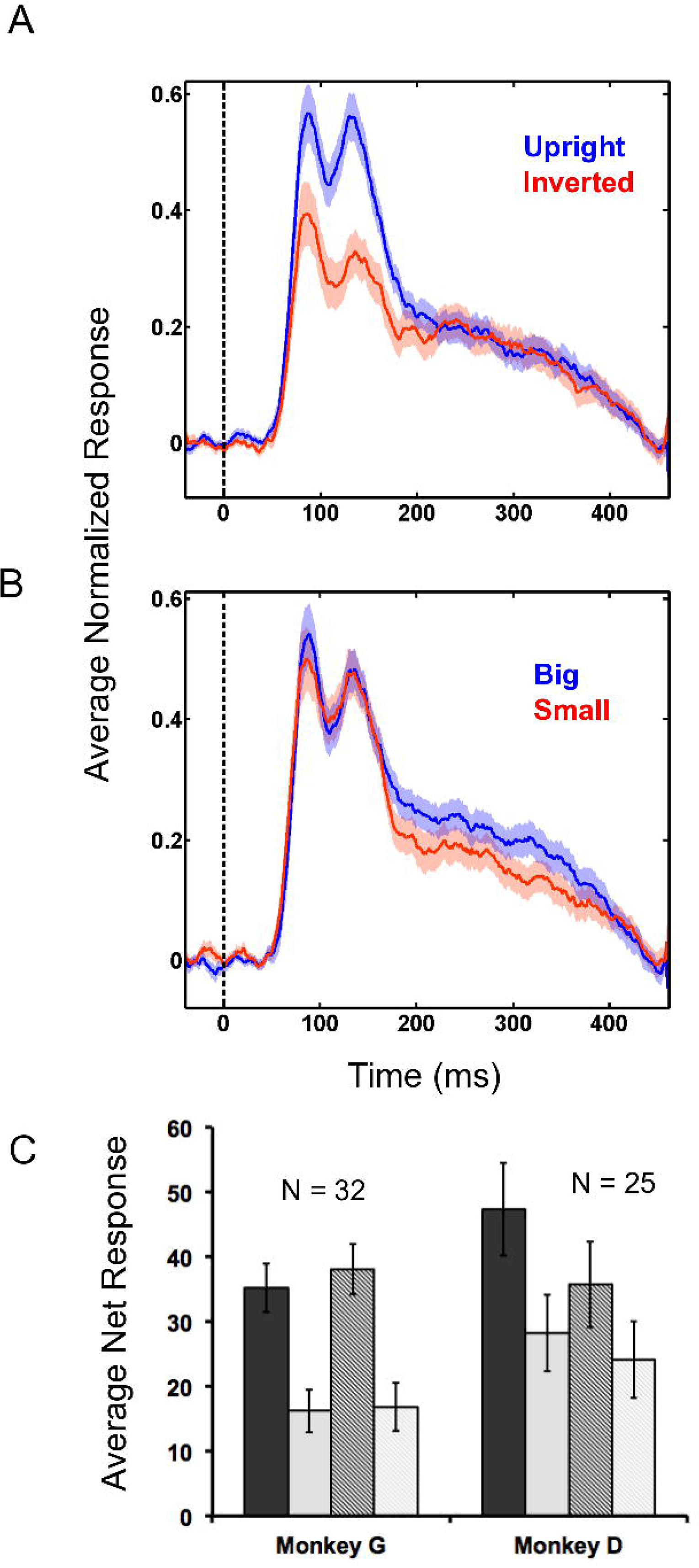
OVERALL ANOVA RESULTS (A) Upright compared to Inverted Faces (averaging across face identity and stimulus size). (B) Large faces compared to Small faces (averaging across face identity and orientation). (C) Individual Monkey Data; All four unique conditions (averaging across face identity) – (from left to right) Upright Large Faces; Inverted Large Faces; Upright Small Faces; Inverted Small Faces.

We found a significant interaction between *Face identity* and *Orientation* implying that picture-plane inversion modulated the average firing rate elicited by the 12 face identities, (F(8.49,475.57) = 4.90, *p* < 0.001; see Figure 3A). This same interaction was significant when the analysis was repeated for each subject separately (i.e. Monkey G, *N* = 32, *p* < 0.01; Monkey D, *N* = 25, *p* < 0.001; See Figure 2C).

**Figure 3:**
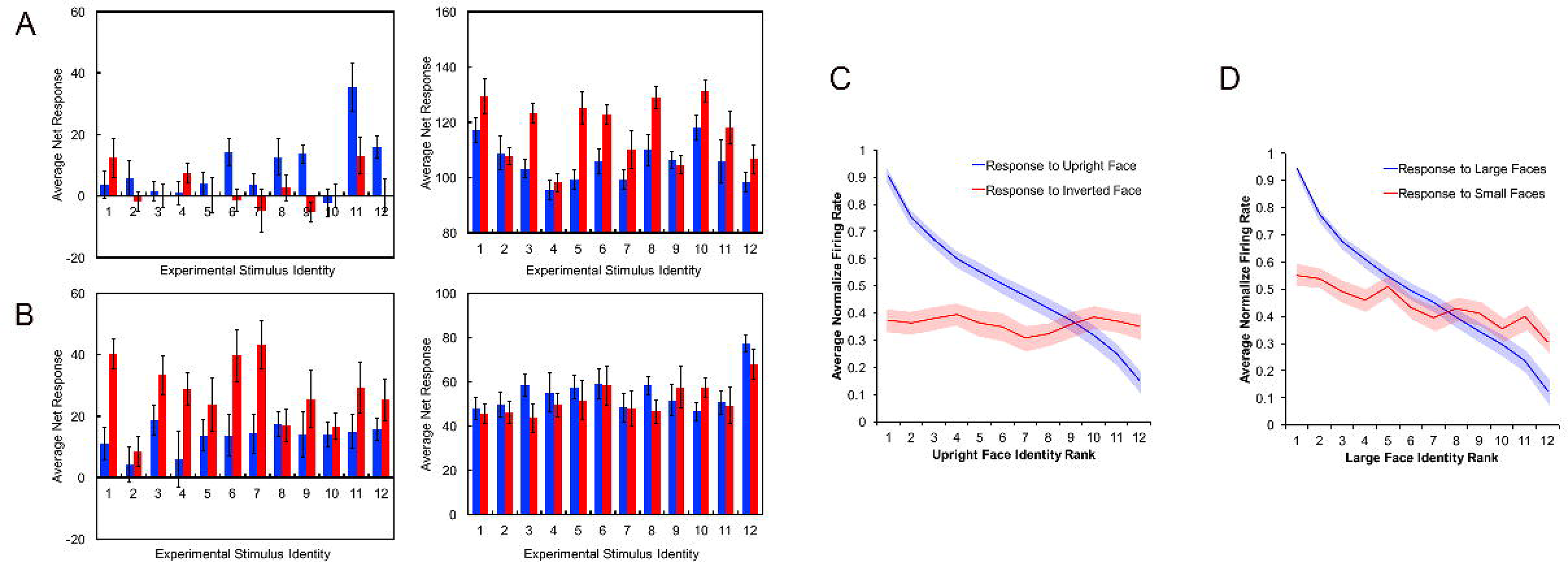
VARIANCE ACROSS NEURONS (A) The average net (raw – baseline) firing rates and standard error (error bars) for 2 randomly selected neurons responding to all 12 face stimuli (blue bars = upright; red bars = inverted). Left was recorded in Monkey D and Right was recorded in Monkey G. (B) The average net (raw – baseline) firing rates and standard error (error bars) for 2 randomly selected neurons responding to all 12 face stimuli (blue bars = large; red bars = small). Left was recorded in Monkey D and Right was recorded in Monkey G. (C) Line graph indicating the average normalized firing rate as a function of stimulus identity, after stimuli have been ranked based on the average response in the upright face condition (i.e. the upright identity that elicited the highest average firing rate was ranked as number 1 and is represented on the far left of the x-axis). (D) Line graph indicating the average normalized firing rate as a function of stimulus identity, after stimuli have been ranked based on the average response in the large face condition (i.e. the upright identity that elicited the highest average firing rate was ranked as number 1 and is represented on the far left of the x-axis).

#### Ranking Analysis of identity preference

To determine whether each neuron’s preference among the upright face stimuli was independent of its preference among the inverted face stimuli, we compared identity preference in the orientation conditions using the rank approach^23,24^. For each neuron, the upright face trials were used for ranking the 12 identities according to response strength in descending order (from best to worst) and the inverted face trials were analysed using the upright face rankings (responses averaged across size). A one-way ANOVA on the ranked inverted face data showed no evidence of a decrease in response as a function of upright face rank (F(8.663,485.14) = 0.53, *p* = 0.88; G-G corrected; see Figure 3C). Thus, these results provide no indication that identity preferences, when averaging across neurons, were the same in the upright and inverted conditions.

We also performed this same ranking analysis on the two size conditions (10 dva hereafter referred to as ‘large’; and 5 dva hereafter referred to as ‘small’). For each neuron, the large face trials were used for ranking the 12 identities according to response strength in descending order (from best to worst) and the inverted face trials were analysed using the upright face rankings. For this analysis, responses were averaged across the manipulation of orientation (upright and inverted trials). A one-way repeated measures ANOVA on the ranked small face data revealed evidence for a preserved rank (F(8.196,458.959) = 3.599, *p* < 0.001; G-G corrected; see Figure 3C). One follow-up test was used to confirm the linear trend (F(1,56) = 21.429, *p* < 0.001).

### Variability across neurons and classifier performance

The above analysis indicates that across the population of face-selective neurons recorded in area ML, there was a difference in the average response to upright and inverted faces. However, the same neurons showed a similar mean response to faces with different presentation sizes. Importantly, the ranking analysis unpacked an important difference between these affine manipulations: stimulus rank was preserved across the size manipulation but not the orientation manipulation.

To further probe whether our images manipulations differentially affected identity decoding in area ML we used the correlation coefficient classifier described in the methods section. The percentage of correct classifications (classifier score) is indicative of how accurate the population of ML neurons can identify an upright or inverted face. First, we classified face identity within the two orientation conditions (ignoring differences in size). The classification score obtained with upright faces was 38.62%, which was above the 99^th^ percentile of the corresponding null distribution (median = 7.5%; 2^nd^ percentile = 2.5%; 98^th^ percentile = 16.67%). For the classifier on inverted face trials performance was 36.51%, which was also above the 99^th^ percentile of the corresponding chance distribution (median = 7.5%; 2^nd^ percentile = 2.5%; 98^th^ percentile = 16.67%). To test whether the classifier score for inverted faces was different from the classifier score for upright faces we, first, created an upright performance distribution by repeating the classifier procedure, with correct identity labels, 1000 times (see Methods). The 5^th^ and 95^th^ percentiles for this upright score distribution were 30.83% and 45.83% respectively. The classification score for inverted faces (i.e. 36.51%) was in the 29^th^ percentile of the upright distribution and, thus, there was no evidence that the inverted score was sampled from a different distribution.

Finally, we tested cross-orientation classifier performance (i.e. training with upright data and testing with inverted and vice versa). The classifier performed more poorly when trained with data from upright face trials and tested with data from inverted face trials (*classifier performance* = 10.10%) which was below the 75^th^ percentile of the null distribution computed by training the classifier with randomly shuffled identity labels (chance distribution median score = 8.3%; 2^nd^ percentile = 3.3%; 98^th^ percentile = 14.17%). Likewise, when we trained the classifier with inverted face trials and tested again upright face data, classifier performance was low (9.19%) and below the 70^th^ percentile of the chance distribution (median score = 8.3%; 2^nd^ percentile = 2.5%; 98^th^ percentile = 15%). Collectively these observations provide no clear evidence that the classifier could accurately decode identity across the orientation conditions at a level greater than chance. These results are consistent with the results of the rank analysis above, which also suggested there was little correspondence between the response pattern to upright and inverted faces.

We then examined classifier performance for identity when data were restricted to the large and small face trials (with orientation trials combined). The classification score for large face trial data was 31.54%, falling above the 99^th^ percentile of the null distribution (median = 8.3%; 2^nd^ percentile = 3.3%; 98^th^ percentile = 15%). The classification score for small face trial data was also significantly above chance (classifier performance = 16.09%, 96^th^ percentile; null distribution median score = 8.3%; 2^nd^ percentile = 2.5% and 98^th^ percentile = 16.67%). Interestingly, when a same cross-size decoding procedure was performed on large and small face trials (i.e. the classifier was trained with large face trials and tested on data from small face trials) the classifier performed at a level above the 95^th^ percentile of the chance distribution obtained by shuffling identity labels. For instance, trained with large face trials and tested with small face trials, classifier performance was 29.55%. This score was above the 99^th^ percentile of the chance distribution (median score = 7.5%; 2^nd^ percentile = 2.5 and 98^th^ percentile = 16.67%). Results were similar when we trained the classifier with data from small face trials and tested against data from large face trials: Classifier performance was 15.95% which was above the 99^th^ percentile of a chance distribution (median score of 8.3%, 2^nd^ percentile = 3.3%, 98^th^ percentile = 14.17%).

## Discussion

Overall, our results obtained by recording in the face-selective middle patch (ML) of the monkey IT confirm that the individuality of a human face picture can be decoded from the firing rates of a modest number of face-selective neurons, i.e. using the output of 57 neurons classifier performance was above chance for all four conditions. The observation that there is a distinct “neural code” for a set of 12 independent and “naturally occurring” human faces is an important replication of previous work^11,13,25^, here sampling face-selective units exclusively in area ML, a functionally defined area of the cortical face processing network in rhesus monkeys. Although we cannot distinguish between norm-based and feature-based coding (see^13,25^), we confirm that the pattern of activity elicited from a relatively small number of ML neurons varies with stimulus identity reliably across trials even without preselecting the preferred identity for any given neuron.

The average response of the neurons we recorded showed size tolerance and, on average, the neurons responded less to inverted faces than to upright faces^3,5,26^. Further, using a ranking analysis, we found evidence that average identity preferences were tolerant of a decrease in stimulus-size but not picture-plane inversion. These observations converge with previous findings to indicate that the average response of face-selective neurons in area ML are more sensitive to changes in stimulus orientation than they are to changes in stimulus size.

While there was no evidence that the cross-orientation classifier performed above chance, the performance of the cross-size classifier indicates that the population response to stimulus identity was tolerant of a change in stimulus size. These findings suggest that identity-selectivity at a population level is dependent on stimulus orientation but not on stimulus size. However, we also observed classifier performance based on inverted trials was well above chance. Moreover, this performance was not significantly lower from classifier performance based on upright orientation. This means that, even though, the average response of the population is reduced, and the identity preferences change when faces are inverted – there is no information loss for inverted faces in area ML. We note that this observation is in agreement with the lack of behavioral inversion effect in macaque monkeys ^27–29^.

While the role of the face-selective area ML in the monkey face processing network remains controversial, it has been suggested that area ML builds representations of face stimuli at a population level that are shape-dependent^13^. Our results are largely consistent with this conclusion, demonstrating that both the average response of a population and the population code are sensitive to changes in stimulus orientation. We also show that the distinct population response in area ML to individual faces is not, simply, sensitive to all affine images transformations; the pattern of responses across 57 neurons to 12 different faces tolerated a change in stimulus size. That is, scaling the faces down to half their original height results in no change in the average response magnitude, and the sampled neurons retained their identity-selectivity.

## Methods

### Subjects and Localization

We used fMRI to localize the face-selective patches in two male monkeys (*Macaca mulatta*), D and G. Animal care and experimental procedures were approved by the ethical committee of the KU Leuven medical school. To optimize the signal-to-noise ratio, we used an iron oxide contrast agent (monocrystalline iron oxide nanoparticle or MION; the details of this procedure are described elsewhere^30^). Eighty images of faces, bodies, fruits, manmade objects and hands (16 images per category) were presented to the monkeys in blocks during continuous fixation. These images have been used to isolate face-selective cells in previous studies of rhesus monkeys^3,15^ and were presented on a square canvas with a height that subtended a visual angle of 8°. Consistent with previous reports, there were several discrete regions (face-selective patches) in both monkeys that responded more to faces than the four other non-face categories. Single unit recordings were performed in three regions in the right hemisphere of both subjects. All recordings were in the lateral lip of the lower bank of the Superior Temporal Sulcus (STS; Figure 1A) in the middle lateral face patch (ML). ML was located ~4mm anterior to the interaural line in monkey D and ~6mm anterior to the interaural line in monkey G (see Figure 1A).

### Single Cell Procedure and Analysis

We surgically implanted a plastic recording chamber in both monkeys that targeted ML and isolated 882 single neurons in total, using epoxylite-insulated tungsten microelectrodes (FHC) and standard electrophysiological procedures described in detail elsewhere^5,24^. Stimuli were displayed on a CRT display (Philips Brilliance 202 P4; 1024 × 768 screen resolution; 75 Hz vertical refresh rate) at a distance of 57 cm from the monkey’s eyes. The 32 images (16 faces, 16 non-face objects) that were used to search for responsive neurons and measure their face-selectivity were taken from the 80 images that were used in the fMRI block-design localizer. The 16 non-face objects were taken from 4 different categories (headless bodies, hands, gadgets, and fruits), selected to be similar to faces in their round shape (e.g. an orange or a closed fist). All images were 8° of visual angle in height, width was allowed to vary. For electrophysiological recordings, however, the noise background was removed from these images and replaced with a uniform grey background and then gamma corrected.

The position of the subject’s right eye was continuously tracked by means of an infrared video-based tracking system (SR Research EyeLink; sampling rate 1KHz). A monkey initiated a trial by fixating on a central fixation spot (size = 0.2° of visual angle) that was always present throughout the trial. The monkey was then required to fixate on this spot (within a 2° x 2° fixation window) for 300 ms prior to stimulus onset and during the stimulus presentation (300 ms). An additional 300 ms fixation period after stimulus offset was required before the monkey was rewarded for continuous fixation with a fluid reward. Trials were separated by an inter-stimulus interval of at least 500 ms, the exact duration being dependent on the oculomotor behavior of the monkey in between the trials (see Figure 1B). In the main tests, the stimuli were at the center of the screen, behind the fixation spot. Each trial presented a monkey with a single stimulus, drawn from the set in a pseudo random order. Each stimulus was repeated at least twice for every neuron discriminated. In each recording session we recorded the first single unit encountered at the predetermined depth with respect to the silence associated with the sulcus, regardless of face-selectivity or visual responsiveness. Each unit thereafter was at least 150μm deeper than the previous.

After a neuron’s spike was isolated, we recorded its response to Face / Non-face stimuli (at least 2 trials per stimuli) in order to compute the neuron’s face-selectivity index. Following previous studies ^3,5,16^, we defined for each neuron a face-selectivity index as FSI = (mean net response faces – mean net response nonface objects)/(| mean net response faces | + | mean net response nonface objects|). We counted a neuron as being “face-selective” if the FSI was greater than 0 (meaning it’s average response to face stimuli was greater than its average response to non-face stimuli).

Without further selection, we then tested each neuron using an independent image set comprised of 12 achromatic faces of unfamiliar adult Caucasian individual (6 females). The height of these stimuli subtended 10° of visual angle in the “large” condition and 5° in the “small” condition. The timing parameters were identical to those described for the category-search procedure. All images depicted neutral expressions and were frontward facing. External cues to facial identity (e.g., hair, ears, and neck) were removed using Adobe Photoshop. The luminance and root-mean square (RMS) contrast of all stimuli were adjusted to match the mean luminance and contrast values of the entire image set. To create the “inverted” stimuli we rotated the 12 “upright” faces 180° in the picture-plane.

Firing rate was computed for each unaborted trial in two analysis windows: a baseline window ranging from 250 to 50 ms before stimulus onset and a response window ranging from 50 to 350 ms after stimulus onset. Responsiveness of each recorded neuron was tested offline by a split-plot ANOVA with repeated measure factor baseline versus response window and between-trial factor stimulus. Only neurons for which either the main effect of the repeated factor or the interaction between the two factors was statistically significant (i.e. *p* < 0.05) were analyzed further. Net firing rate during a trial was calculated by subtracting the firing rate in the baseline window from that in the response window. There were 12 face identities that were presented in each of the four experimental conditions (upright large / upright small / inverted large / inverted small). There were, thus, 48 different conditions in total. These 48 conditions were repeated in at least 5 trials per neuron. In order to pool across neurons and monkeys, we normalized the data with respect to the maximum response across the 48 conditions (averaging across trials) for each neuron, using the net response.

#### Pattern Classifier

To assess face identity coding we used a correlation coefficient classifier^8,31^ based on zero-one loss measure. In this analysis, a pattern classifier is trained on a subset of data to derive the presented stimulus from the pattern of activity in a population of neurons. The proportion of correct classifications (classifier accuracy) is indicative of how well the neural activity pattern represents face identity.

The classifier was trained on the pattern of activity across all 57 neurons on 83.3% of the trials, and tested on the remaining 16.6%. To do this, we first examined classifier performance based on the upright face trials. Excluding trials where the stimulus was presented upside down we used a subset of 6 randomly selected trials per combination of neuron (57 neurons) and face identity (12 identities). For the training, 12 vectors, one for each upright face identity, were created containing the average responses of all 57 neurons on 5 of the 6 selected trials (i.e. 12 vectors of 57 average responses).

For the test phase, a vector with the responses of all 57 neurons on a remaining 6^th^ trial was correlated with each of the 12 vectors created using the training phase. The trained identity that yielded the highest correlation with the test identity was used as the predicted identity (the classifier was correct if the predicted identity was equal to the test identity). This procedure was repeated 6 times, once for each of the selected trials to serve as the test data. To minimize trial selection influence, this total procedure was repeated 100 times, with random permutations of the selection of trials for each neuron. To eliminate the influence of net differences in firing rate of the individual neurons in the correlation, for each permutation, before training and testing, the data were Z-scored (using the mean and standard deviation within each neuron, across orientation and stimuli). The average percentage of ‘correct’ decisions of the classifier was used as the classifier performance.

To test statistical significance of the classifier performance against chance, a null distribution was generated by repeating the above described procedure 1000 times, while, randomly shuffling the identity labels of the stimuli during training. For each of these 1000 repetitions, classifier performance was obtained, resulting in a probability distribution. The proportion of data points from this distribution higher than the performance obtained by using the real face identities indicated the probability level of the classifier not being better than chance (in theory; 1/12 or 8.3%).

## Acknowledgements

We thank Christophe Ulens, Marc De Paep, Sara De Pril, Wouter Depuydt, Astrid Hermans, Piet Kayenbergh, Gerrit Meulemans, Inez Puttemans and Stijn Verstraeten for technical and administrative assistance. The funders had no role in study design, data collection and analysis, decision to publish, or preparation of the manuscript

## Author Contributions Statement

Experiment was designed by all authors. Data collection was performed by J.T. The data were analyzed by J.T and G.V.B. All authors wrote the main manuscript text and prepared the figures. All authors reviewed and approved the manuscript.

